# FORCAST: a fully integrated and open source pipeline to design Cas-mediated mutagenesis experiments

**DOI:** 10.1101/2020.04.21.053090

**Authors:** Hillary Elrick, Viswateja Nelakuditi, Greg Clark, Michael Brudno, Arun K. Ramani, Lauryl M.J. Nutter

**Affiliations:** Centre for Computational Medicine, The Hospital for Sick Children, Toronto, ON M5G 1X8, Canada; The Centre for Phenogenomics, The Hospital for Sick Children, Toronto, ON M5T 3H7, Canada; Department of Computer Science, University of Toronto, Toronto, ON M5T 3A1, Canada; University Health Network, Toronto, ON, Canada

**Author notes:** These authors contributed equally to this work (Hillary Elrick, Viswateja Nelakuditi).

## Abstract

Cas-mediated genome editing has enabled researchers to perform mutagenesis experiments with relative ease. Effective genome editing requires tools for guide RNA selection, off-target prediction, and genotyping assay design. While independent tools exist for these functions, there is still a need for a comprehensive platform to design, view, evaluate, store, and catalogue guides and their associated primers. The Finding Optimizing and Reporting Cas Targets (FORCAST) application integrates existing open source tools such as JBrowse, Primer3, BLAST, bwa, and Silica to create a complete allele design and quality assurance pipeline. FORCAST is a fully integrated software that allows researchers performing Cas-mediated genome editing to generate, visualize, store, and share information related to guides and their associated experimental parameters. It is available from a public GitHub repository and as a Docker image, for ease of installation and portability.

---

With the advent of Cas-mediated genome editing, a wide range of tools have been made available to aid the experimental design process. This includes CRISPOR^1^, GuideScan^2^, and CHOPCHOP^3^ to design and evaluate guides, Cas-OFFinder^4^ to predict off target sites, Benchling (https://benchling.com) to generate and store guides, and the original MIT website for scoring guides^5^ (now offline). While these tools assist Cas-mediated experimental design, they are each tailored to individual parts of this process. Thus, there exists a need for a single free, versatile, and fully integrated software that allows researchers performing Cas-mediated genome editing to generate, visualize, store, and share information related to guides and their associated experimental parameters.

We developed the open-source tool Finding Optimizing and Reporting Cas Targets (FORCAST) to provide such an integrated functionality. FORCAST is available for use with any organism and utilizes rigorous criteria to generate and rank guides, perform quality assurance of alleles, and design genotyping primers. By maintaining an internal database, FORCAST allows users to save and retrieve existing design details as required. Users can also mark existing guides and primers that are no longer being used, based on the results obtained from *in silico, in vitro*, or *in vivo* experiments. Specificity scores, predicted off-target sites, and genomic context can then be used to make an informed decision for primer and guide redesign of a particular target.

All information saved in FORCAST is stored only on local infrastructure, which provides full control of data and eliminates security and privacy concerns. Additionally, the use of Docker allows for deployment in a cloud environment and the leverage of high performance computing when available. Thus, FORCAST acts as a shared resource within a laboratory to prevent duplication of effort and facilitate coordination of Cas-mediated genome editing experiments.

## Results

FORCAST has been used to design and genotype over 176 successful Cas-mediated *in vivo* gene knockout experiments in mice, at the time of writing. It has been extensively tested by model production teams for bugs, and features have been added to improve the workflow. A typical design workflow using FORCAST is described in Figure 1.

**Figure 1.**
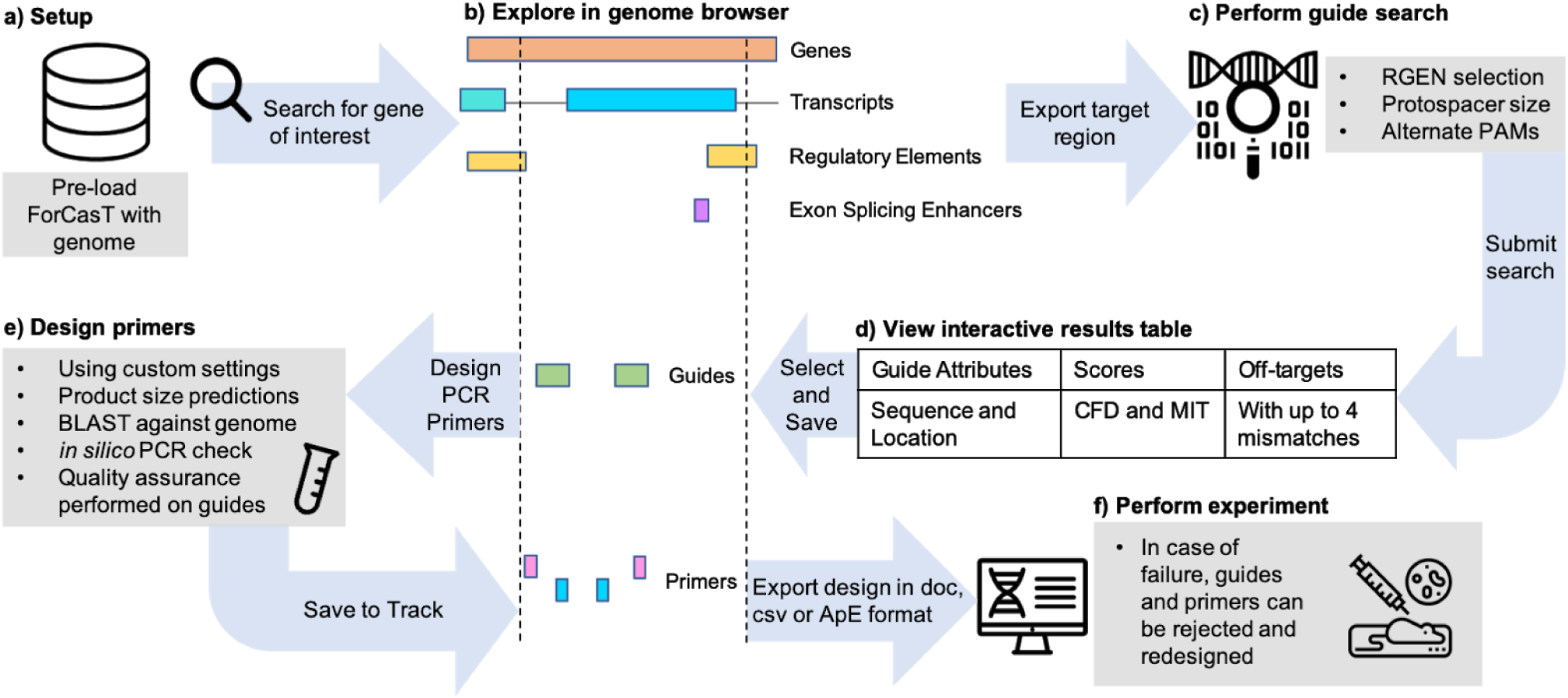
Performing a Cas-mediated experiment with FORCAST. **a)** Any Ensembl genome can be selected for use in the tool. **b)** An interactive genome browser with default tracks (Genes, Transcripts, Regulatory Elements) as well as custom tracks such as Exon Splicing Enhancers allows users to explore their region of interest. **c)** Users refine their guide search by selecting an RNA-guided endonuclease (RGEN), protospacer length, and types of potential off-target sites to consider. **d)** All guides in the search region are displayed in a ranked table with their available scores and potential off-target sites. Saved guides are displayed in the Genome Browser with a user-defined label and notes. **e)** Genotyping primers can be designed for the wild-type (WT) and endonuclease-mediated (EM) alleles. Each potential primer is checked for specificity against the genome and quality assurance is performed on guides to ensure only transcripts of the selected gene are affected within the edited region. **f)** Design details can be exported in various formats. In the case of a design failure, the experiment can be revisited in FORCAST and existing guides or primers rejected and re-designed.

Though *Streptococcus pyogenes* Cas9 (SpCas9)^6^ remains the most commonly used system for genome editing, FORCAST enables researchers to use other RNA-guided endonucleases (RGENs) which provide advantages including reduced size^7^, increased number of target-able sites^8^, and generation of staggered cuts^9,10^, expanding the array of RGEN-mediated genome edits scientists are able to make. FORCAST comes preloaded with the following RGENs: *Streptococcus pyogenes* Cas9 (SpCas9), *Acidaminococcus* Cas12a (AsCpf1/Cas12a), *Streptococcus canis* Cas9 (ScCas9), and *Staphylococcus aureus* Cas9 (SaCas9). Relevant information about these RGENs, described in Table 1 below, is loaded into the application at setup. New RGENs can be easily added to the database, and the default specifications can be modified as needed. Novel RGENs with genome editing capabilities are being discovered at a rapid pace, and FORCAST was designed so that researchers can quickly use these new technologies as they emerge.

**Table 1:** RNA-guided endonuclease (RGEN) attributes. Specific attributes for common RGENs loaded into FORCAST include protospacer adjacent motifs (PAMs), sequence length of the guide RNA, RGEN cleavage site, spacer seed region, known non-canonical PAMs, and any available implemented scores.

| Common Name | Protospacer Adjacent Motif (PAM) | Guide RNA Spacer Sequence Length | Cleavage Site | Seed | Published non-canonical PAMs | Implemented Scores |
| --- | --- | --- | --- | --- | --- | --- |
| SpCas9 | NGG, 3' end<sup>11</sup> | 17-20bp<sup>12</sup> | 3bp upstream from 3' end<sup>13</sup> | 12bp from 3' end<sup>14</sup> | NAG, NCG, NGA<sup>15</sup> | MIT<sup>5</sup>, CFD<sup>22</sup> |
| AsCpf1/Cas12a | TTTV, 5' end<sup>9</sup> | 20-23bp<sup>9</sup> | 19bp from 5' end on + strand, 23 bp from 5' end on - strand<sup>10</sup> | 6bp from 5' end<sup>10</sup> | TTTT, CTTA, TCTA, TTCA<sup>10</sup> | - |
| ScCas9 | NNG, 3' end<sup>8</sup> | 20bp<sup>8</sup> | 3bp upstream from 3' end<sup>8</sup> | 12bp from 3' end<sup>8</sup> | - | - |
| ScCas9 | NNGT, 3' end<sup>8</sup> | 20bp<sup>8</sup> | 3bp upstream from 3' end<sup>8</sup> | 12bp from 3' end<sup>8</sup> | - | - |
| SaCas9 | NNGRRT, 3' end<sup>7</sup> | 21-23bp<sup>7</sup> | 3bp upstream of PAM<sup>7</sup> | 7-8bp from 3' end<sup>7</sup> | NNGRR<sup>7</sup> | - |

### Performance

Benchmark tests were performed to compare FORCAST’s guide searching speed and accuracy to CRISPOR, GuideScan, and Cas-OFFinder, three frequently used tools in guide design (Table 2). Command-line versions of each tool were tested by an automated program (Supplementary Data) on a server with 8GB of RAM using randomly selected input search sequences.

**Table 2:** Benchmarking of FORCAST, CRISPOR, GuideScan, and Cas-OFFinder.

| **Table 2: Benchmarking of FORCAST, CRISPOR, GuideScan, and Cas-OFFinder.** |  |  |  |  |
| --- | --- | --- | --- | --- |
| **Tool** | **FORCAST** | **CRISPOR** | **GuideScan** | **Cas-OFFinder** |
| 150 bp region (chr14:54165316-54165466) |  |  |  |  |
| Avg. Time Elapsed* (seconds) | 19.63 | 26.04 | 0.36** | 693.34 |
| Guides Returned | 21 | 21 | 17 | (21)*** |
| Potential Off-Target Sites Identified | 12172 | 6377 | 399 | 14657 |
| 300 bp region (chr6:136920211-136920511) |  |  |  |  |
| Avg. Time Elapsed (seconds) | 28.50 | 28.02 | 0.75 | 1206.75 |
| Guides Returned | 49 | 49 | 47 | (49)*** |
| Potential Off-Target Sites Identified | 21820 | 12329 | 875 | 26973 |
| 600 bp region (chr10:20497604-20498204) |  |  |  |  |
| Avg. Time Elapsed (seconds) | 37.29 | 31.21 | 0.73 | 1486.82 |
| Guides Returned | 64 | 64 | 42 | (64)*** |
| Potential Off-Target Sites Identified | 23864 | 12563 | 841 | 2665443 |
| 1200 bp region (chr5:74093060-74094260) |  |  |  |  |
| Avg. Time Elapsed (seconds) | 63.35 | 62.61 | 0.81 | 3624.55 |
| Guides Returned | 184 | 184 | 131 | (184)*** |
| Potential Off-Target Sites Identified | 48356 | 33673 | 1734 | 73670 |
| * Each tool was tested three times on the input sequence; we report the results averaged across tests.<br>** GuideScan precomputes a database of gRNAs for a particular RGEN and genome.<br>*** Cas-OFFinder only provides functionality to search for off-targets; guides were supplied to the tool. |  |  |  |  |

FORCAST is significantly faster than Cas-OFFinder and returns many more off-targets sites than GuideScan. It is comparable in speed to CRISPOR while still returning more off-target sites. Accurately reporting the number of potential off-target sites is essential for reducing the risk of undesired edits, calculating scores, and performing quality control on resulting organisms. Though GuideScan has the fastest running time, it is limited to showing only off-target sites with three mismatches in the genome. CasOFFinder was set to return off-targets with four mismatches for this test, though it allows for up to nine. FORCAST and CRISPOR return off-targets with up to four mismatches, as off-target sites with up to four mismatches have been shown to produce undesired edits^16^. Furthermore, FORCAST reports potential off-targets adjacent to non-canonical PAMs (NAG, NCG, and NGA for SpCas9), with the option to modify this list. To increase speed, FORCAST processes and displays a maximum of 1000 potential off-target sites for a given guide, and skips scoring guides in repetitive regions by default. However, these restrictions can be disabled by users to display a full list of guides and their off-targets for a given region. With these options, FORCAST allows users to decide whether to prioritize speed or completeness when searching for and evaluating guides and off-targets.

Furthermore, we tested FORCAST in a region of the genome (chr19:10907072-10907187) that several tools were reported to erroneously suggest guides with a high number of potential off-target sites^17^. Rather than rejecting these guides outright, FORCAST displays a warning about the high number of mismatches and reports them at the bottom of the ranked results table.

## Discussion

### Advantages

FORCAST is available as an open source stand-alone application (see Availability), which provides several benefits over publicly accessible web or cloud-based tools, such as security, privacy, and long-term data storage and integrity. Data saved by the tool is stored only on local or owned cloud infrastructure, giving organizations full control of their data including the ability to backup, export and share experiment details. These qualities make FORCAST ideal for use in a Core Facility, where standard protocols and controlled access to data is essential. FORCAST can also be used with Docker, making it suitable for non-technical users to run on a personal computer with minimal setup.

Additionally, we recognized that laboratories have specific needs and protocols with regard to experimental design and validation. Flexibility was kept in mind during the development of FORCAST, allowing researchers to use specific genome versions, modify available RGENs, include additional annotation data, and define custom primer design settings. This makes FORCAST an incredibly flexible tool that can be used to aid in the design of Cas-mediated genome editing experiments across various fields of biology.

### Future Directions

FORCAST is under active development and planned features include adding the ability to design conditional alleles, mutations in non-coding genes, and point mutations (variants). Additional goals include incorporating genomic variant information associated with a reference genome, and incorporating new scoring methods, including specificity scores for AsCas12a.

## Methods

### Implementation

Several open-source tools are integrated into FORCAST; these tools were chosen for their demonstrated reliability, accuracy, and ease of use. JBrowse^18^ is used to visualize genomic features such as genes, transcripts, regulatory information, guides, and primers. BWA^19^, BEDtools^20^, and SAMtools^21^ are used to find guides and their potential off-target sites. The published MIT^5^ and Cutting Frequency Determination^22^ (CFD) scores are used to evaluate guide specificity. While Primer3^23^ is used to generate PCR primers for genotyping, BLAST^24^ and Silica (https://www.gear-genomics.com/silica) are used to evaluate primer specificity. FORCAST is written in Python and uses MongoDB to store genes, guide RNA spacer sequences with their associated RGENs, and PCR primers for quality control and genotyping.

Detailed installation and setup instructions for FORCAST are described in the GitHub repository (see <u>Availability</u>). Briefly, a shell script installs all required tools and programs, after which users can populate FORCAST with their genomes of interest using the included Python setup script. During setup, the genome sequence (in FASTA format) and genomic annotation (in GFF3 format) files are downloaded programmatically from the Ensembl FTP site. Users can specify the version of Ensembl release to use; if this isn’t specified, the latest version is used. A BED file categorizing the genome into intergenic, intronic, and exonic regions is generated from the annotation file and gene symbols, identifiers, and chromosomal locations are extracted and stored in the MongoDB. Genome sequence and annotation files are then loaded into JBrowse. Additionally, BLAST and BWA indexes are built at setup, for which we recommend at least 8GB of RAM.

### Availability

Project home page: https://ccmbioinfo.github.io/FORCAST

Demo: https://youtu.be/SJMDAuJRuDI

Operating systems(s): Host machine must be Docker-compatible (most Linux distributions, MacOS 10.12 and higher, Windows 10) or run Ubuntu 16.04 to host natively. Web-interface is operating system-independent, tested on Chrome, Firefox, and Opera.

Programming languages: Python, JavaScript, bash

Other requirements: Recommended that host machine has at minimum 8GB of RAM License: GPLv3 License

## Supporting information

Supplemental Benchmarking Code

## Acknowledgements

Thanks to Lauri Lintott at The Centre for Phenogenomics for beta testing and feedback as well as beta testers Denise Lanza, Jason Heaney, Juan Gallegos, Kiran Rajaya, and Vivek Ramanathan at Baylor College of Medicine. Thanks to Mia Husic for constructive feedback and proofreading. Thank you also to Tobias Rausch for assistance integrating the *in silico* PCR tool Silica. This work was funded by Genome Canada and Ontario Genomics (OGI-137) and supported by the Canadian Centre for Computational Genomics (C3G), part of the Genome Innovation Network (GIN), funded by Genome Canada through Genome Quebec and Ontario Genomics.

