## Supplemental Benchmarking Code for "FORCAST: a fully integrated and open source pipeline to design Cas-mediated mutagenesis experiments": compareTools.py

```
#!/bin/python3

import sys
import os
import re
import subprocess
import time
from subprocess import PIPE, DEVNULL
from randomLocation import *

def call_guideScan(searchInput, gene):
  switch_version = subprocess.call(["pyenv","local", "2.7.12/envs/guidescan"])
  if switch_version == 0:
    time_before = time.time()
    p = subprocess.Popen(["guidescan_guidequery","-b","tools/guidescan_public/guidescan-source_code/cas9_mm10_all_guides.bam","-c", searchInput,"--target=within","-o=output/guidescan"], stdout=PIPE)
    out, err = p.communicate()
    time_after = time.time() - time_before
    if out.decode('utf-8') != '':
       print(out)
    if err:
      print(err)
  else:
    sys.exit("Unable to switch to pyenv for guidescan")

  # check to ensure file was written successfully
  output_file = searchInput+"_within.csv"
  if not os.path.exists(os.path.join("output/guidescan", output_file)):
    print("Output file for "+str(searchInput)+" not found")
    return False, 0
  
  return True, time_after

def call_forcast(searchInput, gene):
  time_before = time.time()
  p = subprocess.Popen(["python3","/var/www/html/CasCADe/src/guide-finder/GuideSearchAndScore.py", "--genome=mm10", "--gene="+str(gene), "--input="+str(searchInput),"--output=output/forcast/"+str(searchInput)+".csv"], stdout=PIPE)
  out, err = p.communicate()
  time_after = time.time() - time_before
  if err:
    print(err)
    return False, 0

  return True, time_after

def call_casOFFinder(searchInput, gene):
  """ first need to identify guides using own modules and write to file which is used as input """
  # use ForCasT code to get the sequence/gRNAs for input region
  forcast_dir = "/var/www/html/CasCADe"
  output_fasta = "output/tempfiles/"+str(searchInput)+".fa"
  p = subprocess.Popen(["python3",
    os.path.join(forcast_dir,"src/guide-finder/core/get_sequence.py"), 
    os.path.join(forcast_dir,"bin/twoBitToFa"), searchInput, 
    os.path.join(forcast_dir,"jbrowse/data.mm10/downloads/mm10.2bit"), output_fasta], stdout=PIPE
  )
  out, err = p.communicate()
  if err:
    print(err)  
  
  # use fasta file of sequence to find gRNAs
  sys.path.append(os.path.join(forcast_dir,"src/guide-finder/core"))
  from find_grna import find_grna
  result = find_grna('1','20',output_fasta)
  # put the results into a file and run/time CasOFF
  casoff_input = 'output/tempfiles/'+str(searchInput)+".input"
  with open(casoff_input, 'w') as casoff_file:
    casoff_file.write("/var/www/html/CasCADe/jbrowse/data.mm10/downloads/mm10.fa\n")
    casoff_file.write("NNNNNNNNNNNNNNNNNNNNNRG\n")
    for guideID, guide in result.items():
      casoff_file.write(guide['guide_seq']+"NNN 4\n")
  
  casoff_output = 'output/casoffinder/'+str(searchInput)+".csv"
  time_before = time.time()
  p = subprocess.Popen(["./tools/cas-offinder", casoff_input, "C", casoff_output], stdout=PIPE)
  out, err = p.communicate()
  time_after = time.time() - time_before
  
  if err:
    print(err)
    return False, 0
 
  return True, time_after

def call_crispor(searchInput, gene):
  search_fasta = "output/tempfiles/"+str(searchInput)+".fa"
  if not os.path.exists(search_fasta):
    # use ForCasT code to get the sequence for input region
    forcast_dir = "/var/www/html/CasCADe"
    output_fasta = "output/tempfiles/"+str(searchInput)+".fa"
    # create the input fasta if it hasn't been created already
    create_input = subprocess.Popen(["python3",
    os.path.join(forcast_dir,"src/guide-finder/core/get_sequence.py"), 
    os.path.join(forcast_dir,"bin/twoBitToFa"), searchInput, 
    os.path.join(forcast_dir,"jbrowse/data.mm10/downloads/mm10.2bit"), output_fasta], stdout=DEVNULL
    )
    out, err = create_input.communicate()
    if err:
      print(err)

  switch_version = subprocess.call(["pyenv","local", "crispor2"])
  if switch_version == 0:
    # now run the command
    time_before = time.time()
    output_file = "output/crispor/"+str(searchInput)+".csv"
    run_crispor = subprocess.Popen(["./tools/crisporWebsite/crispor.py","mm10",search_fasta,output_file], stdout=PIPE, stderr=DEVNULL)
    out, err = run_crispor.communicate()
    time_after = time.time() - time_before
    print(out)
    if err:
      print(err)
      return False, 0
    return True, time_after
  else:
    print("unable to switch python version")
    return False, 0

def count_guidescan(searchInput):
  """ counts the number of guides returned and their off-targets"""
  results_file = "output/guidescan/"+str(searchInput)+"_within.csv"
  g_count = 0
  ot_count = 0
  if os.path.exists(results_file):
    f = open(results_file, "r")
    for i in range(0,3):
      f.readline() # remove first 3 lines from file
    for line in f:
      g_count += 1
      sum_ot = line.split(",")[7]
      #print(sum_ot)
      ot_count += int(sum_ot)
      
  return g_count, ot_count

def count_forcast(searchInput):
  """ return the number of guides and off targets in the searchInput """
  results_file = "output/forcast/"+str(searchInput)+".csv"
  g_count = 0
  ot_count = 0
  if os.path.exists(results_file):
    f = open(results_file, "r")
    for line in f:
      found_line = re.match(r"(\d*)\s*GUIDES\sFOUND", line)
      offtg_line = re.match(r"(\d*)\s*OFF\sTARGETS\sFOUND", line)
      if found_line:
        g_count = int(found_line.group(1))
      elif offtg_line:
        ot_count += int(offtg_line.group(1))
  
  return g_count, ot_count

def count_casOFFinder(searchInput):
  casoff_output = 'output/casoffinder/'+str(searchInput)+".csv"
  ot_count = 0
  guides_seen = []
  if os.path.exists(casoff_output):
    f = open(casoff_output, "r")
    for line in f:
      ot_count += 1
      guide = line.split("\t")[0]
      if guide not in guides_seen:
        guides_seen.append(guide)

  return len(guides_seen), ot_count

def count_crispor(searchInput):
  results_file = "output/crispor/"+str(searchInput)+".csv"
  guide_count = 0
  off_target_count = 0
  if os.path.exists(results_file):
    f = open(results_file, "r")
    f.readline() # remove header
    for line in f:
      guide_count += 1
      off_target_count += int(line.split("\t")[5])

  return guide_count, off_target_count

def repeatTests(TOOLS, n=150, m=5):
  # repeat the test for selected tools on m randomly generated regions of size n
  gene, searchInput = selectRandomRegion('mm10',n)

  for i in range(0,m):
    tool_rand=list(TOOLS.items()) # shuffle tools so they're run in random order
    random.shuffle(tool_rand)
    for tool, opts in tool_rand:
      if opts['run']:
        print(tool+str(" RUNNING..."))
        time_before = time.time()
        result, time_elapsed = opts['call_function'](searchInput, gene)
        opts['time_elapsed'].append(time_elapsed)
        opts['success'] = result
        if result:
          print(tool+str(" FINISHED."))
          print(str(time_elapsed)+" seconds elapsed.")
          g_count, ot_count = opts['count_function'](searchInput)
          opts['guide_count'] = g_count
          opts['offtarget_count'] = ot_count
          print("Guides found: "+str(g_count))
          print("Off-targets found: "+str(ot_count))
  
  for tool, opts in TOOLS.items():
    if opts['run']:
      print(tool + " average time elapsed: "+ str(sum(opts['time_elapsed'])/len(opts['time_elapsed'])))
      print(str(opts['guide_count']) + " guides found")
      print(str(opts['offtarget_count']) + " off targets reported")

def main():
  TOOLS = {
    'GUIDESCAN': {
      'run': True,
      'success': False,
      'time_elapsed': [],
      'call_function': call_guideScan,
      'count_function': count_guidescan
    },
    'FORCAST': {
      'run': True,
      'success': False,
      'time_elapsed': [],
      'call_function': call_forcast,
      'count_function': count_forcast
    },
    'CASOFFINDER': {
      'run': True,
      'success': False,
      'time_elapsed': [],
      'call_function': call_casOFFinder,
      'count_function': count_casOFFinder
    },
    'CRISPOR': {
      'run': True,
      'success': False,
      'time_elapsed': [],
      'call_function': call_crispor,
      'count_function': count_crispor
    }
  }
  repeatTests(TOOLS)

if __name__ == "__main__":
  main()
```
