## Supplemental Benchmarking Code for "FORCAST: a fully integrated and open source pipeline to design Cas-mediated mutagenesis experiments": randomLocation.py

```
#/bin/python3

import json
import random
import sys
from pymongo import MongoClient

""" 
REQUIRES A FORCAST SET UP DATABASE FOR THE GIVEN GENOME TO BE RUNNING ON THE SERVER
"""

def selectRandomRegion(genome, length):
  # select a random gene from the given organism and return a region of the specified length within the gene
  try:
    # connect to the db
    client = MongoClient()
    db = client[genome]
  except Exception as e:
    print("Error fetching genome for: "+str(genome))

  # get most recent ensembl release
  curr_release = -1
  for c in db.collection_names():
    if 'geneInfo' in c:
      if int(c.split("_",1)[1]) > int(curr_release):
        gene_collection = db[c]
        curr_release = int(c.split("_",1)[1])

  # fetch random gene from collection
  rand_gene = None
  # re-retrive until non-predicted gene found
  while not rand_gene or rand_gene['Name'].startswith("Gm") or rand_gene['Name'].endswith("Rik"):
    rand_gene = list(gene_collection.aggregate([{"$match": {}}, {"$sample": {"size": 1}}]))[0]
  # get random location in gene range
  coord = random.randrange(int(rand_gene['start']), int(rand_gene['end']), int(length))

  # return random location
  gene = rand_gene['Name']
  loc = str(rand_gene['chr'])+":"+str(coord)+"-"+str(int(coord)+int(length))
  print(gene + " " + loc)
  return gene, loc

def main():
  # parse command-line args
  if len(sys.argv) != 3:
    print("Please provide a genome and desired length of random (within a gene) coordinates")
    sys.exit()

  selectRandomRegion(sys.argv[1], sys.argv[2])

if __name__ == "__main__":
  main()
```
